# Engineering lithoheterotrophy in an obligate chemolithoautotrophic Fe(II) oxidizing bacterium

**DOI:** 10.1101/2020.09.30.321166

**Authors:** Abhiney Jain, Jeffrey A. Gralnick

## Abstract

Neutrophilic Fe(II) oxidizing bacteria like *Mariprofundus ferrooxydans* are obligate chemolithoautotrophic bacteria that play an important role in the biogeochemical cycling of iron and other elements in multiple environments. These bacteria generally exhibit a singular metabolic mode of growth which prohibits comparative “omics” studies. Furthermore, these bacteria are considered non-amenable to classical genetic methods due to low cell densities, the inability to form colonies on solid medium, and production of copious amounts of insoluble iron oxyhydroxides as their metabolic byproduct. Consequently, the functional understanding of these bacteria remains speculative despite the availability of substantial genomic information. Here we develop the first genetic system in neutrophilic Fe(II) oxidizing bacterium and use it to engineer lithoheterotrophy in *M. ferrooxydans*, a metabolism that has been speculated but not experimentally validated. Our work suggests that *M. ferroxydans* partitions energy generation from carbon oxidation. This synthetic biology approach could be extended to gain physiological understanding and domesticate other bacteria that grow using a single metabolic mode.

## Introduction

Among its many biological roles, Fe(II) is an important energy source in the environment as an electron donor for chemolithoautotrophic growth. Diverse neutrophilic chemolithoautotrophic bacteria have been known to oxidize Fe(II) in many circumneutral environments^1–6^. Understanding the metabolism and physiology of chemolithoautotrophic Fe(II) oxidizing bacteria is of environmental and ecological importance because of their widespread impact in various environments on multiple biogeochemical cycles including iron, carbon, nitrogen, phosphorous and other metals^1–8^. While a substantial amount of genomic information about chemolithoautotrophic Fe(II) oxidizing bacteria exists, functional knowledge remains speculative in the absence of metabolic and genetic studies. Chemolithoautotrophic Fe(II) oxidizing bacteria grow at aerobic-anaerobic interfaces^1–8^ where they outcompete chemical oxidation of Fe(II) by oxygen. The requirement for this specific niche makes these organisms difficult to culture for laboratory studies. These bacteria are generally metabolic specialists which grow by a singular metabolic mode of oxidizing Fe(II) as the energy source while fixing carbon dioxide and respiring low levels of oxygen^1–8^. The single metabolic growth mode prohibits comparative “omics” studies to gain a deeper functional understanding of these microorganisms. Furthermore, chemolithoautotrophic Fe(II) oxidizing bacteria are considered non-amenable to genetic methods because these bacteria do not grow on solid medium, produce low growth yields (10^6^-10^7^ cells/mL) and accumulate a substantial amount of insoluble iron oxyhydroxide as their obligate metabolic byproduct^1–8^. The inability to grow and form colonies on solid medium prevents the application of traditional genetic methods to select and screen for mutants. Growth by a single metabolic mode prohibits targeted gene deletions to probe Fe(II) oxidation and carbon flow as mutants defective in these pathways will be unable to grow. Low cell yield along with the presence of insoluble iron oxyhydroxide presents a formidable challenge for DNA transformation and subsequent phenotypic analysis to readily test genetic parts and methods.

Here we use synthetic biology to study *Mariprofundus ferrooxydans*, the founding member of the *Zetaproteobacteria*^3^ which are thought to be the dominant Fe(II) oxidizers in marine environments^1,8^. We develop genetic methods and tools to transform *M. ferrooxydans* and manipulate its metabolic capacity by expressing foreign genes, yielding an engineered variant capable of using glucose as a carbon source instead of CO_2_.

## Results and Discussion

We developed a conjugation protocol to successfully transform *M. ferrooxydans* using the donor strain *Escherichia coli* WM3064, which is auxotrophic for diaminopimelic acid (DAP)^9^. *M. ferrooxydans* transformed with pRK2m3^10^ continued to grow and produce characteristic twisted iron oxide stalks^2^ over successive transfers in the presence of kanamycin (Fig. 1A, B) while wild-type incubated with kanamycin was unable to grow (data not shown) or produce twisted iron stalks (Fig. 1C). 16S rRNA gene sequencing confirmed that the transformed culture was *M. ferrooxydans*. After ten transfers, *E. coli* cells were undetectable by microscopy or growth in lysogeny broth (LB) medium augmented with DAP. Maintenance of pRK2m3 in the transformed *M. ferrooxydans* cells was confirmed by amplifying plasmid specific DNA using total extracted DNA as the template (Fig. 1D) and verified by sequencing. These results demonstrate that *M. ferrooxydans* was able to replicate pRK2m3 over repeated transfers under kanamycin selection. With a method for transformation and selection established, we were able to express green fluorescent protein (GFP), encoded by *gfpmut2*^11^ and driven by the P_neo_ promoter amplified from upstream the gene encoding kanamycin resistance on pRK2m3. P_neo_ was used because it provided sufficient expression to confer kanamycin resistance in *M. ferrooxydans* transformed with the pRK2m3 vector. Microscopy confirmed production of GFP in the engineered strain (Fig. 2).

**Fig. 1.**
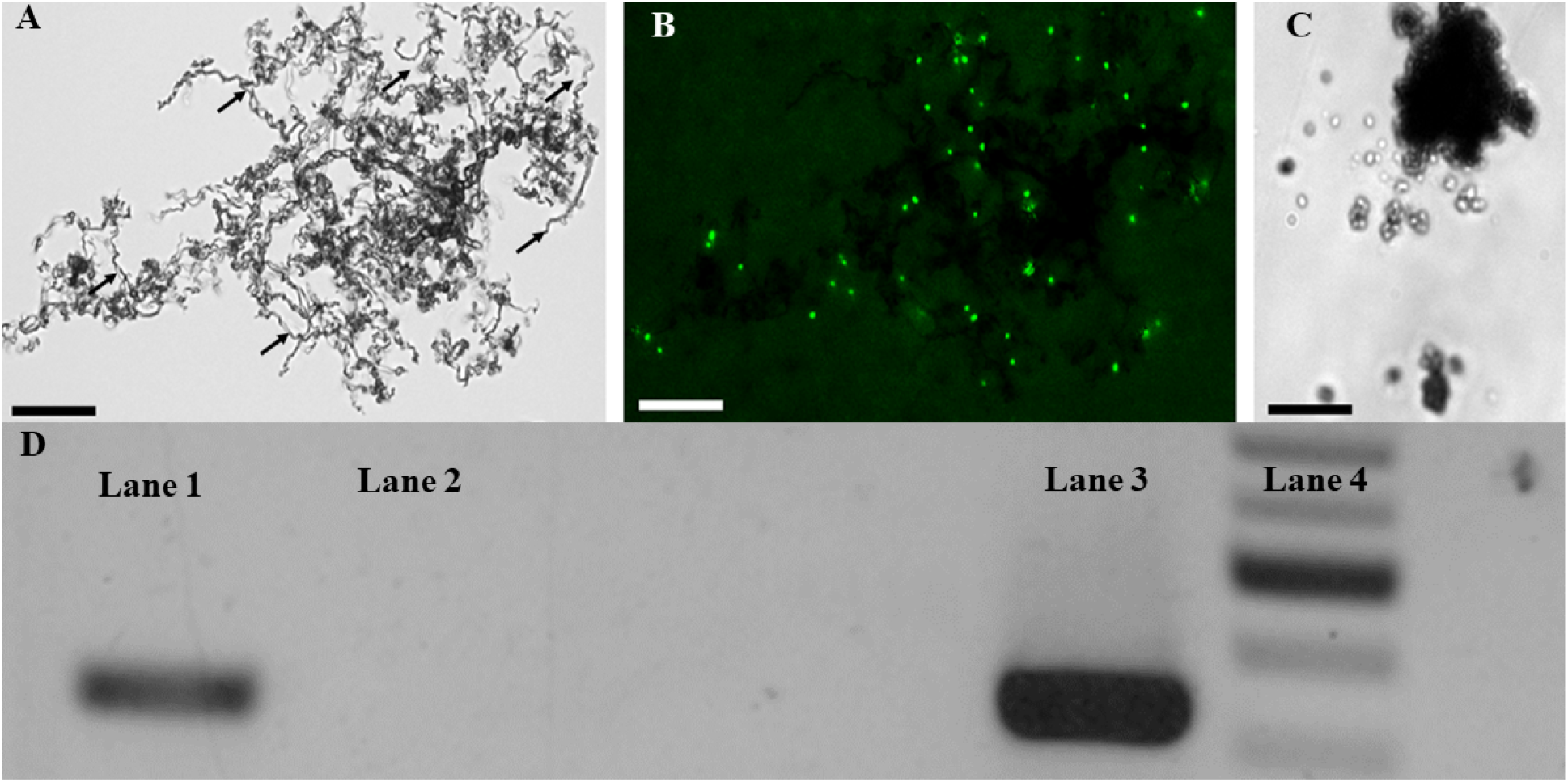
Transformation of *M. ferrooxydans*. Successful transformation of pRK2m3 (empty vector) into *M. ferrooxydans* was confirmed by (A) growth in the presence of 200 μg/mL kanamycin as shown by the production of characteristic stalk formation (black arrows) in the bright field micrograph and (B) an epifluorescent micrograph of the same field showing cells stained with Syto 9. (C) Wild-type cells were unable to grow or produce stalks in the presence of 200 μg/mL kanamycin. Scale bars indicate 25 μm. (D) Electrophoregram showing amplification of pRK2m3 DNA fragments using DNA extracted from the transformed cells (Lane 1) and purified pRK2m3 (Lane 3) as the template. Template DNA containing DNA extracted from the wild-type cells did not produce any amplification (Lane 2). Lane 4 is a DNA ladder.

**Fig. 2.**
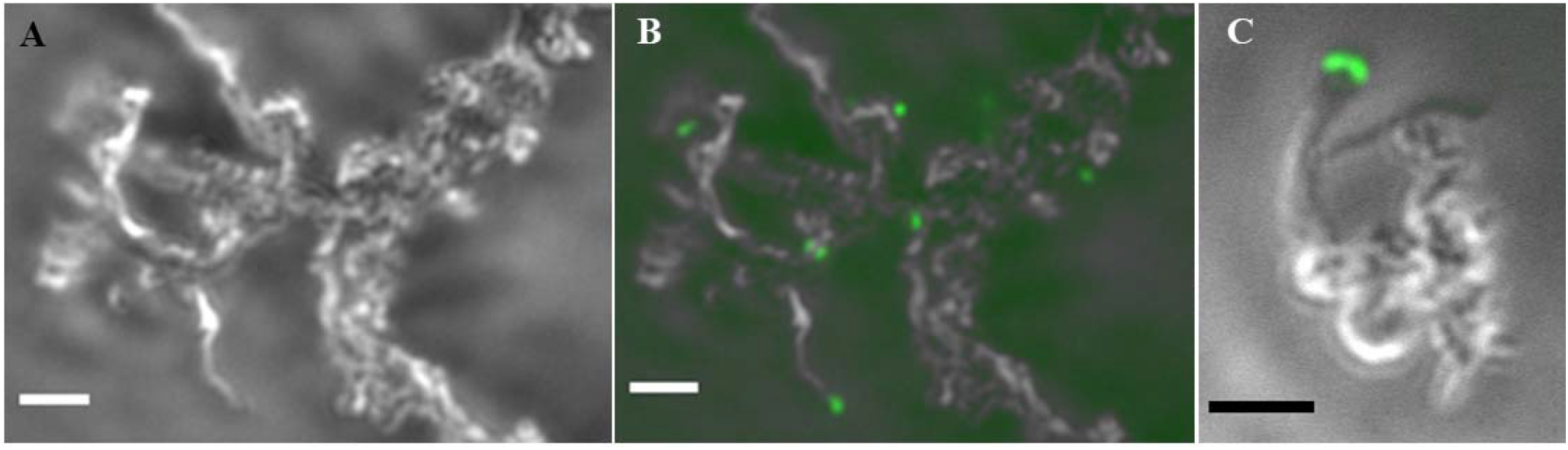
Expression of the green fluorescent protein in *M. ferrooxydans*. (A) Light micrograph showing the characteristic twisted stalks produced by *M. ferrooxydans* containing the plasmid with *gfpmut2* and grown in the presence of 200 μg/mL kanamycin. (B, C) Composite images of light and epifluorescent micrographs showing the green fluorescent cells attached to the stalks. Scale bars indicate 5 μm.

We next sought to leverage our ability to introduce and express foreign genes to augment the metabolism of *M. ferrooxydans*. The *M. ferrooxydans* genome is predicted to encode genes for glycolysis and the Krebs cycle^12^, but lacks genes encoding glucokinase or an apparent glucose transporter. We hypothesized that by introducing the capability to transport and phosphorylate glucose, *M. ferrooxydans* could use it as a carbon and energy source. The genes *galP* and *glk* from *E. coli*, encoding a glucose symporter and glucokinase^13^, were cloned into pRK2m3 to create pGlu where each gene was individually driven by P_neo_ promoters. *M. ferrooxydans* cells transformed with pGlu did not yield viable cells when selected heterotrophically using glucose (data not shown). However, growth was observed when *M. ferrooxydans* cells transformed with pGlu were selected lithoheterotrophically under Fe(II) oxidizing conditions with glucose as the sole carbon source without addition of carbon dioxide (Fig. 3). *M. ferrooxydans* transformed with an empty pRK2m3 vector was unable to grow with glucose as the sole carbon source (Fig. 3). The presence of pGlu was confirmed by amplifying and sequencing *galP, glk* and the kanamycin resistance cassette using total DNA extracted from transformed glucose-grown cells as template (data not shown). Purity of the transformed cells was confirmed by 16S rRNA gene sequencing, microscopy analysis and the absence of bacterial growth in LB medium augmented with DAP.

**Fig. 3.**
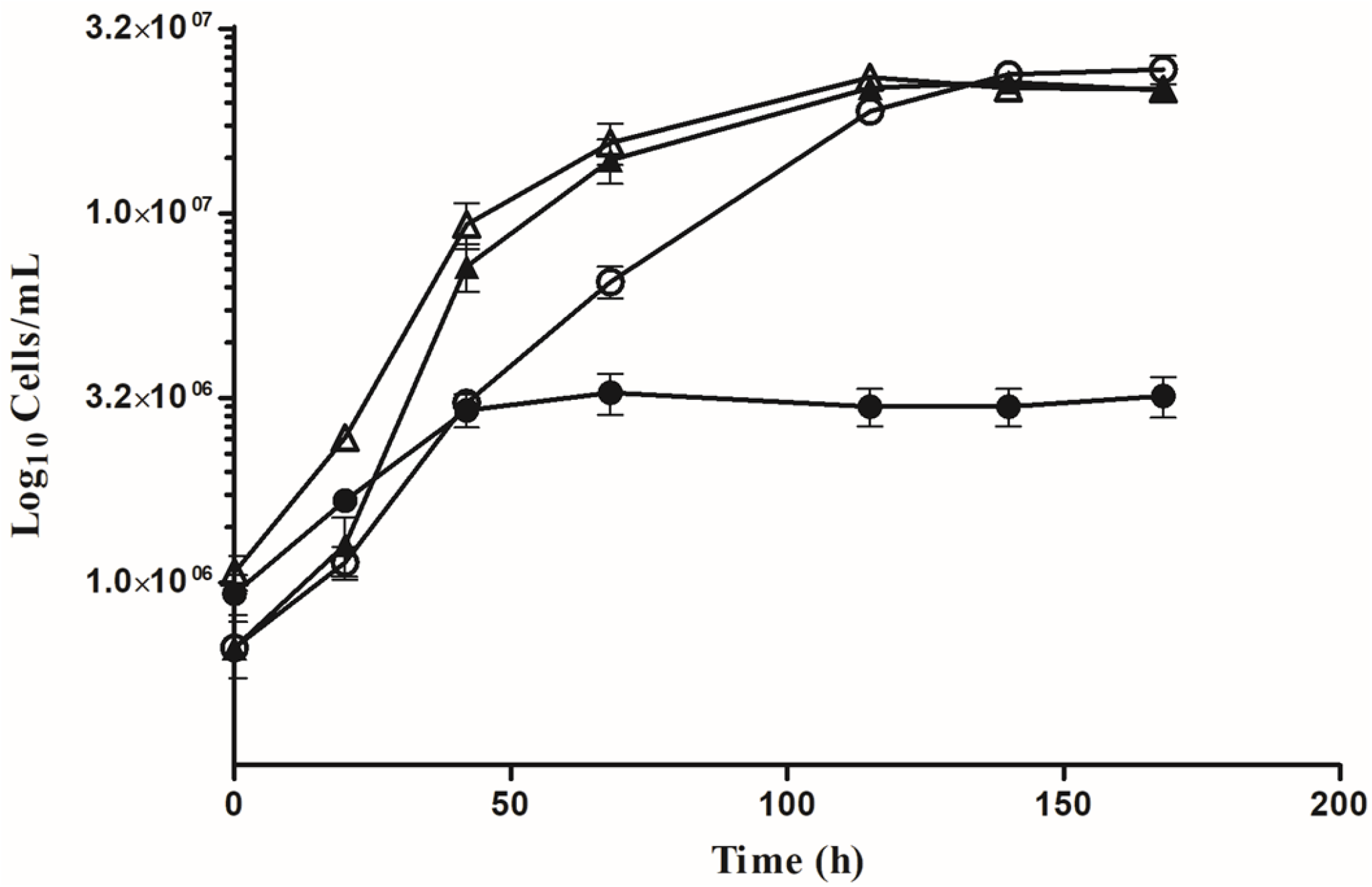
Lithoheterotrophic growth of *M. ferrooxydans* containing pGlu. Growth curves of *M. ferrooxydans* containing pGlu (open symbols) and *M. ferrooxydans* containing empty vector pRK2m3 (closed symbols) grown on either glucose (circles) or carbon dioxide (triangles) as the sole carbon source. Error bars represent standard deviation of three replicates.

The rate of Fe(II) oxidation by the engineered lithoheterotrophic strain was slower during glucose-dependent growth compared to the Fe(II) oxidation rate during carbon dioxide-dependent growth (Fig. 4). Interestingly, the engineered strain oxidized less total Fe(II) when grown with glucose compared to carbon dioxide (Fig. 4), despite achieving similar final cell densities (Fig. 3). The increase in cell yield per unit Fe(II) oxidized during growth on glucose of the engineered strain can be theoretically attributed to additional energy production from glycolysis and/or biomass precursors provided by glucose.

**Fig. 4.**
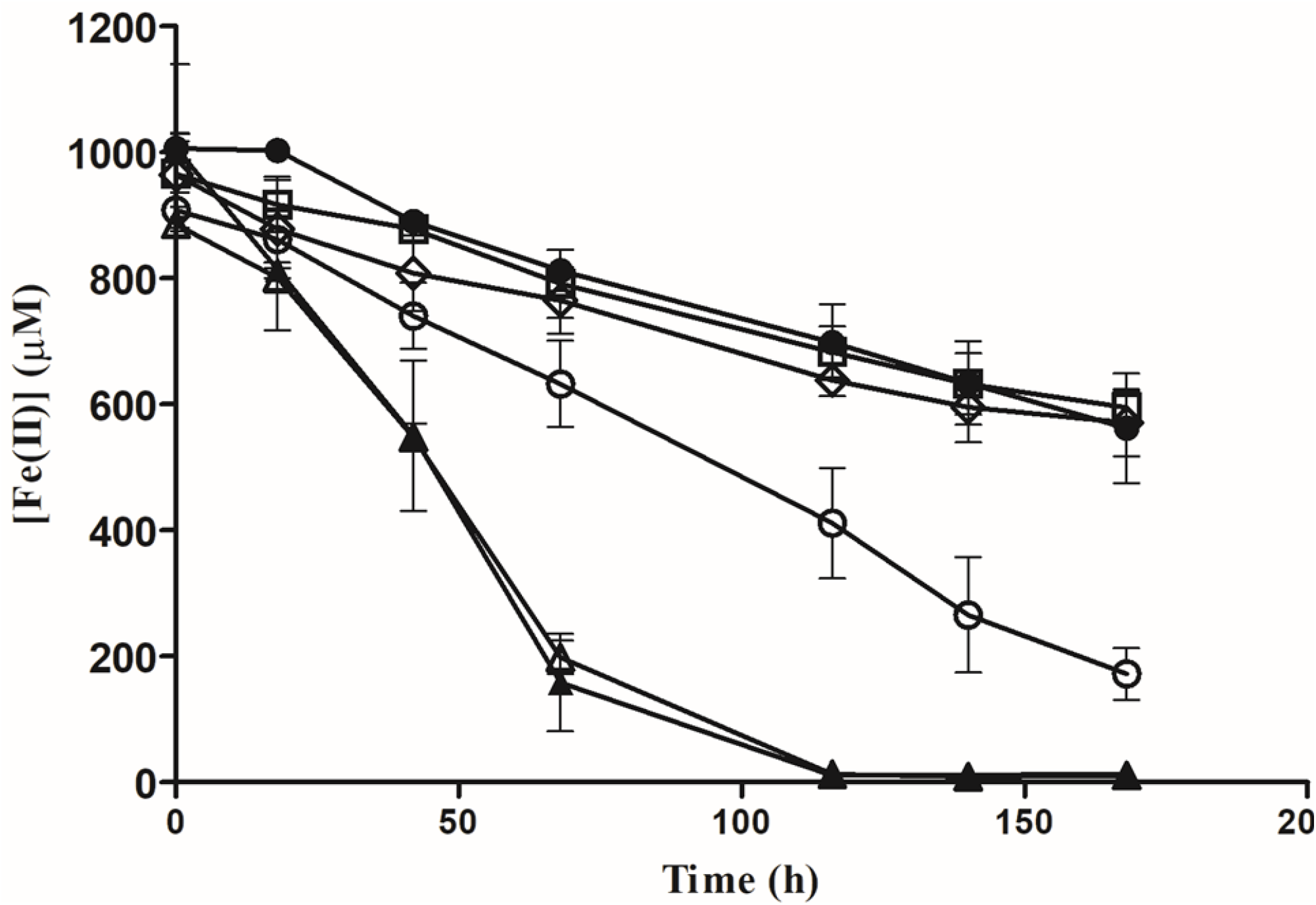
Fe(II) oxidation rate of *M. ferrooxydans* containing pGlu. Fe(II) quantification over time for *M. ferrooxydans* containing pGlu (open symbols) and *M. ferrooxydans* containing empty vector pRK2m3 (closed symbols) grown on either glucose (○) or carbon dioxide (Δ) as the sole carbon source. Fe(II) was also quantified over time for abiotic treatments in the absence of *M. ferrooxydans* cells, containing either glucose (◊) or carbon dioxide (□) in the medium. Error bars represent standard deviation of three replicates.

The inability to transport organic carbon or glycolytic lesions have previously been hypothesized as the reasons for obligate autotrophy in some organisms^14^. However, when these deficiencies were addressed using pGlu in *M. ferrooxydans*, heterotrophic growth was not observed, suggesting that the cells either have insufficient flux through glycolysis or are unable to convert NADH/NADPH produced by glycolysis into proton motive force (and then ATP). The obligate requirement of Fe(II) as the energy source even while using glucose in the engineered lithoheterotrophic strain provides an important insight into the metabolic functioning of *Mariprofundus* where glycolysis seems to be partitioned from energy metabolism. Such a metabolism could be one of the reasons driving obligate lithotrophy in *M. ferrooxydans*. We hypothesize that additional components and alteration of metabolic networks will be required to achieve heterotrophic growth in *M. ferrooxydans*, and possibly other obligate chemolithoautotrophs. Although chemolithoheterotrophy, where Fe(II) oxidation provides energy and organic carbon serves primarily as a carbon source, has been speculated in Fe(II) oxidizing bacteria^15^, it has not been experimentally validated. Our ability to successfully engineer chemolithoheterotrophy in *M. ferrooxydans* suggests that other microorganisms in the environment may also be capable of this growth strategy.

Our work provides a proof of concept for using synthetic biology to augment metabolism in recalcitrant microbes to enhance their growth capabilities in laboratory conditions. This approach may be applied to a wide range of bacteria that live by only a single metabolic mode. Enhancing the metabolic capabilities of metabolic specialists can provide a way to better understand their physiology and provide a blueprint for their domestication.

## Methods

### Bacterial cultivation and DNA extraction

*M. ferrooxydans* was obtained from the National Center for Marine Algae and Microbiota at Bigelow Laboratory and grown in sealed containers containing artificial sea water medium (ASW)^2^. After autoclaving, filtered air was added to the headspace to achieve approximately 1% oxygen concentration as the electron acceptor. Either Fe(0) or FeCl_2_ (400 μM) was added as electron donor. For CO_2_ dependent growth, ASW medium was sparged with N_2_:CO_2_ (80:20). For lithohetrotrophic growth, ASW medium lacking bicarbonate, sparged with argon and augmented with 500 μM glucose was used. DNA was extracted from 40 mL of *M. ferrooxydans* culture using a Qiagen DNeasy PowerSoil kit. *E. coli* WM3064 containing the desired plasmid was grown in LB medium containing 360 μM DAP and 50 μg/mL kanamycin.

### Transformation of *M. ferrooxydans*

A 750 mL culture of *M. ferrooxydans* grown using FeCl_2_ was centrifuged at 1500 × rcf for 3 minutes. The supernatant was then collected and centrifuged at 16000 × rcf for 10 minutes. The pellet obtained was washed with ASW-LB medium (9:1 mixture of ASW and LB) and resuspended in 900 μL ASW-LB. One mL of *E. coli* donor strain culture containing approximately 10^9^ cells was washed with LB medium and resuspended in 100 μL LB. Donor and recipient cells were mixed and centrifuged at 16000 x rcf for 10 minutes, supernatant removed and 5 μL DAP (360 mM) added to the pellet. After incubation at 30°C for 18 hours, the pellet was washed with ASW medium and transformed cells were selected under iron-oxidizing conditions with 200 μg/mL kanamycin (without DAP), while diluting out untransformed *M. ferrooxydans* and *E. coli* cells over successive transfers (each at 1:100 dilution).

### Cell and Fe(II) quantification

Fe(II) was quantified using the ferrozine assay^16^. Cells were fixed in 0.8% paraformaldehyde for 2 hours, stained with 12.5 mM Syto9 and counted using a Petroff-Hausser counting chamber on an epifluorescent microscope.

### Plasmid construction

pRK2m3 specific DNA fragment was amplified using the following primers-GAGCTTATCGGCCAGCCTC and TGTAAAACGACGGCCAGT, P_neo_ was amplified from pRK2m3 using pneoF (GATAGAATTCTTGAGACGTTGATCGGCACG) and pneoR (TAGACTCGAGAACACCCCTTGTATTACTGTTTATGTAAGC) primers. To construct the plasmid for GFP expression, *gfp*mut2^11^ was amplified from pUA66^11^ using gfpF (ACGACTCGAGATGAGTAAAGGAGAAGAACTTTTCACTGGA) and gfpR (TAGAGAGCTCTTATTTGTACAATTCATCCATACCATGGGTA) primers and cloned into pRK2m3 with the P_neo_ cassette driving its expression. To construct pGlu, *galP* and *glk* were amplified from *E. coli* K-12 using galPF (ATTTACTAGTATGCCTGACGCTAAACAGG) / galPR (ATTCGAGCTCTTAATCGTGAGCGCCTATTTCG) and glkF (ACGACTCGAGATGACAAAGTATGCATTAGTCGGT) /glkR (TAGAGAATTCTTACAGAATGTGACCTAAGGTCTG) primers respectively. Amplified *galP* and *glk* were cloned under the control of separate P_neo_ promoters in pRK2m3. All the plasmids were transformed into chemically competent *E. coli* WM3064 [7] cells, followed by selection on LB plates containing 50 μM kanamycin and 360 μM DAP.

## Data availability

Raw cell count and Fe(II) quantification data is available from the corresponding author on reasonable request.

## Acknowledgments

This research was supported by the National Science Foundation Center for Dark Energy Biosphere Investigations (OCE-0939564) post-doctoral fellowship to AJ; and Office of Naval Research grant N00014-13-10552 and National Science Foundation grant MCB-1815584 to JAG. This is C-DEBI contribution XXX (to be determined).

## Author contributions

AJ and JAG designed research; AJ performed research; AJ and JAG analyzed data and wrote the paper.

## Competing interests

The authors declare no competing interests.

## Notes

### Competing Interest Statement

The authors have declared no competing interest.

